# RIF1-Long promotes G1 phase 53BP1 nuclear bodies to protect against replication stress

**DOI:** 10.1101/859199

**Authors:** Lotte P. Watts, Toyoaki Natsume, Yuichiro Saito, Javier Garzón, Masato T. Kanemaki, Shin-ichiro Hiraga, Anne D. Donaldson

## Abstract

Human cells lacking RIF1 are highly sensitive to replication inhibitors, but the reasons for this sensitivity have been enigmatic. Here we show that RIF1 must be present both during replication stress and in the ensuing recovery period to promote cell survival. Of two isoforms produced by alternative splicing, we find that RIF1-Long alone can protect cells against replication inhibition, but RIF1-Short is incapable of mediating protection. Consistent with this isoform-specific role, RIF1-Long is required to promote the formation of the 53BP1 nuclear bodies that protect unrepaired damage sites in the G1 phase following replication stress. Overall, our observations show that RIF1 is needed at several cell cycle stages after replication insult, with the RIF1-Long isoform playing a specific role during the ensuing G1 phase in damage site protection.

## INTRODUCTION

The RIF1 protein has emerged as a central regulator of chromosome maintenance, acting in double strand break repair and DNA replication control (Escribano-Díaz et al., 2013; Chapman et al., 2013; Hiraga et al., 2017). During double strand break repair, RIF1 is recruited by 53BP1, dependent upon phosphorylation of 53BP1 by ATM (Chapman et al., 2013; Virgilio et al., 2013; Escribano-Díaz et al., 2013). Together RIF1 and 53BP1 recruit Shieldin and suppress BRCA1 recruitment to damage sites, opposing homologous recombination-based repair and favouring Non-Homologous End Joining (Bunting et al., 2010; Setiaputra and Durocher, 2019).

RIF1 is also implicated in protecting cells from replication stress (Mazouzi et al., 2016; Buonomo et al., 2009; Kumar and Cheok, 2014). Replication stress can be induced by various conditions including drugs such as Aphidicolin, which interrupts replication fork progression by inhibiting the replicative DNA polymerases alpha, delta, and epsilon (Syvaoja et al., 1990). Replication stress leads to genomic instability, mutation and eventually disease (Ogi et al., 2012; Kerzendorfer et al., 2013; Burrell et al., 2013), so understanding the cellular response is central for understanding accurate genome duplication and the action of replication inhibitors as anti-cancer drugs (Feng et al., 2003; Imai et al., 2016). RIF1-deficient cells are acutely sensitive to replication stress, in fact appearing to be more sensitive to replication inhibitors than to DSB-inducing agents (Buonomo et al., 2009), suggesting that protection from stress is a critical RIF1 function.

Several roles have been described for RIF1 in replication control. RIF1 acts as a Protein Phosphatase 1 (PP1) ‘substrate-targeting subunit’ (Peti et al., 2013) that suppresses replication origin initiation by directing PP1 to dephosphorylate the MCM replicative helicase complex (Davé et al., 2014; Hiraga et al., 2014, 2017; Mattarocci et al., 2014; Alver et al., 2017). RIF1 moreover stimulates origin licensing during G1 phase, and protects replication forks from unscheduled degradation (Hiraga et al., 2017; Garzón et al., 2019; Chaudhuri et al., 2016). However, whether deficiency in these functions accounts for the replication stress sensitivity of cells lacking RIF1 has remained unclear (Feng et al., 2003; Buonomo et al., 2009). RIF1 also acts in mitosis to maintain genomic stability. During anaphase RIF1 is recruited to ultrafine bridges (UFBs), along with the BLM and PICH proteins that ensure proper chromosome segregation (Hengeveld et al., 2015). UFBs are believed to correspond to stretches of underreplicated DNA that escape checkpoint surveillance and persist into mitosis (Bergoglio et al., 2013; Bhowmick et al., 2016). Unresolved DNA damage that passes to daughter cells causes formation of 53BP1 nuclear bodies during G1 phase, which protect the damaged DNA (Bruhn et al., 2014; Lukas et al., 2011; Moreno et al., 2016). RIF1 has also recently been described as functioning at the midbody during cytokinesis (Bhowmick et al., 2019).

The human RIF1 transcript undergoes alternative splicing producing two protein isoforms: a long variant of 2,472 amino acids (‘RIF1-L’), and a short variant (‘RIF1-S’) which lacks 26 amino acids close to the C-terminus of the protein (Xu and Blackburn, 2004). RIF1-S was reported to be more abundant in various cancer cell lines (Xu and Blackburn, 2004), hinting at distinct effects of the isoforms. Although RIF1-L was designated the canonical form, studies using cloned RIF1 have invariably used RIF1-S (Xu and Blackburn, 2004; Xu et al., 2010; Escribano-Díaz et al., 2013; Batenburg et al., 2017), without testing for distinct functions of the isoforms.

## RESULTS & DISCUSSION

### Analysing fluorescent degron-tagged RIF1 reveals highly dynamic cell cycle localisation

We aimed to understand how RIF1 guards against replication stress. First we confirmed in a colony formation assay (CFA) that HEK293 cells depleted for RIF1 are sensitive to the polymerase inhibitor Aphidicolin (Fig. 1A,B). HCT116 cells deleted for both copies of *RIF1* were also sensitive (Fig. 1C). These results imply a specific role for RIF1 in protecting cells under replication stress conditions.

**Figure 1:**
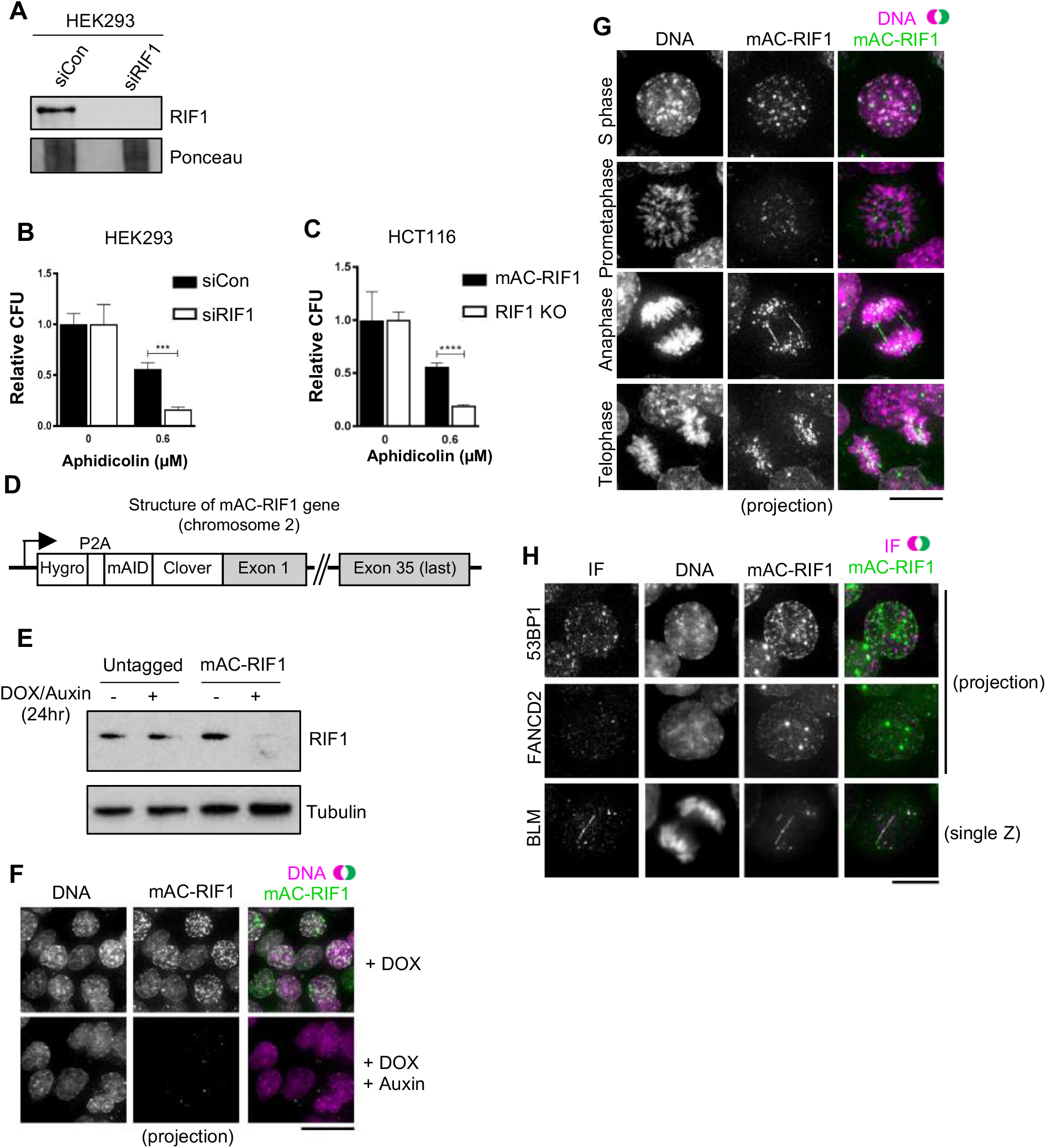
Characterisation of HCT116-based cell lines with Auxin Inducible Degron-tagged RIF1. **(A)** Confirmation of siRIF1 efficacy by western blotting three days after siRNA transfection. Whole cell protein extracts analysed by western blotting with anti-RIF1 and anti-tubulin antibodies. **(B)** Colony Formation Assay (CFA) testing Aphidicolin sensitivity of HEK293 cells treated with siRIF1. Plots show averages and standard deviations of technical triplicates in this and all further figures. ***p<0.001. **(C)** CFA testing Aphidicolin sensitivity of HCT116 mAC-RIF1 and HCT116 RIF1-KO cells. ****p<0.0001. **(D)** Structure of Auxin Inducible Degron (AID)-tagged RIF1 construct (mAC-RIF1), located at both the endogenous RIF1 loci in HCT116 cells carrying the auxin-responsive F-box protein *Oryza sativa* TIR1 (OsTIR1) under DOX control. The RIF1 gene is fused to a tag containing a self-cleaving hygromycin resistance marker, mini-Auxin Inducible Degron (mAID) and monomer Clover (mClover) protein. **(E)** Confirmation of mAC-RIF1 protein degradation. Cells were incubated with 2 μg/ml DOX and 500 μM Auxin for 24 hr, then protein extracts analysed by western blotting. **(F)** mAC-RIF1 degradation assessed by microscopy. mAC-RIF1 cells were treated with 2 μg/ml DOX and 500 μM Auxin for 24 hr. DNA was stained with SiR-DNA and livecell imaging performed using a Deltavision microscope. Scale bar = 10 μm. **(G)** Examples of mAC-RIF1 localisation at different cell cycle stages. DNA was stained with SiR-DNA, and live-cell imaging performed using a Deltavision microscope. Scale bar = 10 μm. **(H)** mAC-RIF1 co-localises with BLM at UFBs but not with 53BP1 or FANCD2 repair proteins. Fixed cells were stained with the above mentioned antibodies and imaging performed using a Deltavision microscope. Scale bar = 10 μm.

Since RIF1 functions at various cell cycle stages, we explored when RIF1 is needed to maintain cell proliferation following replication stress. Specifically, we tested if RIF1 function is required during DNA replication stress, after its occurrence, or both during and after stress. Using Auxin Inducible Degron (AID) technology we constructed a cell line allowing rapid depletion and re-expression of RIF1 at different phases of the cell cycle (Natsume et al., 2016; Nishimura et al., 2009). In an HCT116-based cell line carrying the auxin-responsive degron recognition protein OsTIR1 under DOX control, we N-terminally tagged both RIF1 copies with a degron-Clover construct, termed ‘mAC’, consisting of a mini-Auxin Inducible Degron and monomer Clover (a derivative of GFP (Fig. 1D) (Yesbolatova et al., 2019). The expressed construct remains under control of the endogenous RIF1 promoter. Western blot analysis indicated that expression levels of mAC-RIF1 in the absence of Auxin were similar to those of endogenous untagged RIF1 (Fig. 1E). Treatment for 24 hr with DOX and Auxin led to near-complete degradation of RIF1, as visualised by western blotting (Fig. 1E) and microscopy (Fig. 1F). Using flow cytometry analysis of the RIF1-fused mClover tag, we established minimum concentrations of DOX and Auxin to allow effective depletion (Fig. S1A,B). Degradation was largely complete after 3 hr of Auxin treatment (Fig. S1C), while expression was restored to almost normal levels 5 hr after Auxin removal (Fig. S1D). Together, these results confirm the construction of a cell line allowing rapid depletion and re-expression of RIF1.

We visualised mAC-RIF1 based on Clover fluorescence in the first study of the dynamic behaviour of tagged endogenous RIF1 in living cells. In live-cell imaging experiments we observed a pattern of numerous mAC-RIF1 foci throughout S phase nuclei (Fig. 1G, top row), often with 3-6 prominent foci superimposed on a pattern of more numerous smaller foci, consistent with previous studies (Xu and Blackburn, 2004; Buonomo et al., 2009; Yamazaki et al., 2012). In prometaphase mAC-RIF1 exhibits a novel, circumpolar localisation pattern similar to that described for kinetochores (Fig. 1G, second row, showing polar view of condensed chromosomes) (Magidson et al., 2015). At anaphase, even in unperturbed cells, mAC-RIF1 was frequently at structures apparently corresponding to ultrafine bridges (UFBs) (Fig. 1G, third row). These structures were not stained by conventional DNA dyes (Fig. 1G, third row) but colocalized with BLM, a marker for UFBs (Fig. 1H, bottom panel) (Chan et al., 2007; Barefield and Karlseder, 2012; Hengeveld et al., 2015). At telophase, mAC-RIF1 formed multiple small foci associated with separated chromosomes (Fig. 1G, bottom row), as described previously (Yamazaki et al., 2012; Xu and Blackburn, 2004). Time-lapse imaging of HCT116 mAC-RIF1 cells containing an mCherry-tagged PCNA revealed intense RIF1 foci that accumulated through S phase and G2 (Video 1), but disappeared by metaphase (Video 2). RIF1 signal was absent for a short period at metaphase, quickly followed by reappearance of numerous smaller foci on telophase chromosomes coupled with localisation at anaphase bridges.

The mAC-RIF1 fusion therefore retains the localisation and functional characteristics of the endogenous RIF1 protein. Its highly dynamic behaviour as revealed by this live-cell study indicates that RIF1 functions in several cell cycle phases to maintain chromosome stability.

### RIF1 is needed during and after replication stress to promote cell proliferation

Prolonged auxin-induced degradation of RIF1 caused sensitivity to Aphidicolin, comparable to an HCT116 RIF1-knockout cell line (RIF1 KO) (Fig. 2A, Fig. S1E, Fig. 3B lane 4). To investigate when RIF1 function is required to protect against the effects of Aphidicolin, we synchronised cells first in G1 phase with Lovastatin (Rao et al., 1999; Moghadam-Kamrani and Keyomarsi, 2008), and 8 hr after release from Lovastatin added Aphidicolin to induce replication stress (Fig. 2B, upper timeline). The reversible CDK1/cyclin B1 inhibitor RO-3306 (Vassilev et al., 2006) was simultaneously added, to induce a temporary G2 arrest and prevent cells from proceeding into mitosis. After 28 hr, Aphidicolin and RO-3306 were removed to allow release, and then 4 hr later cells were plated for CFA measurement of cell viability (Vassilev et al., 2006). Synchronisation timings were optimised using flow cytometry analysis of cell cycle progression (Fig. 2C, Fig. S1F).

**Figure 2:**
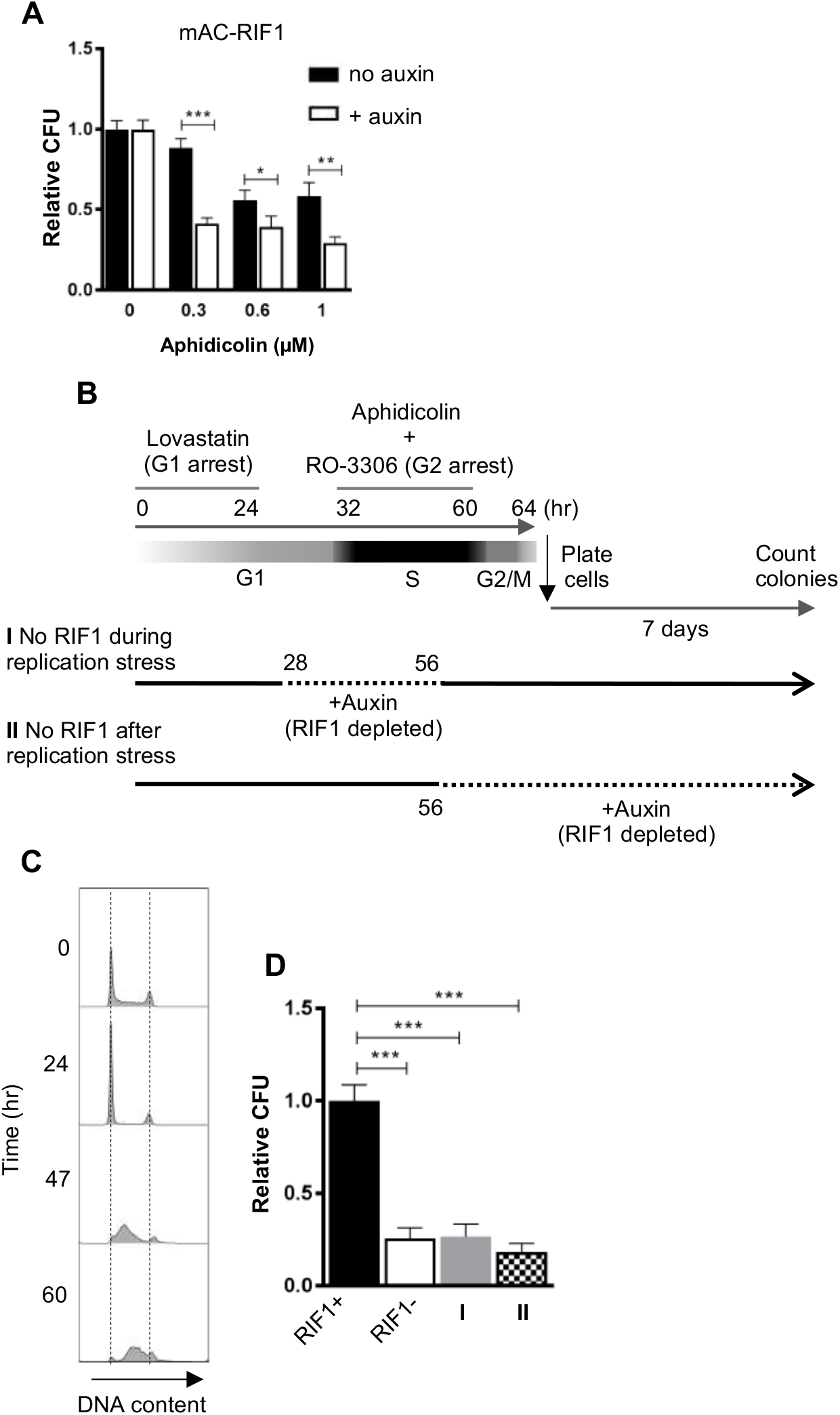
RIF1 is essential during and after drug treatment to protect against effects of replication stress. **(A)** CFA testing Aphidicolin sensitivity of HCT116 mAC-RIF1 cells depleted for RIF1. Cells were treated with DOX and 500 μM Auxin for 48 hr before seeding at low density and treatment with Aphidicolin concentrations indicated. ‘No Auxin’ cells were treated with DOX and Auxinole. *p<0.05; **p<0.01; ***p<0.001 **(B)** Procedure for testing effect of depleting RIF1 during or after Aphidicolin treatment in synchronised cultures. HCT116 mAC-RIF1 cells were incubated with Lovastatin to arrest cells in G1. After 24 hr, cells were released from G1 arrest with Mevalonic Acid. 4 hr into release (28 hr), RIF1 depletion was induced in ‘condition I’ cells by addition of DOX and Auxin. 8 hr into release (32 hr), 1 μM Aphidicolin was added (to both cultures) to induce replication stress and simultaneously, RO-3306 was added to arrest cells in G2 phase. 24 hr later (at 56 hr), in ‘condition I’ cells RIF1 was re-expressed by removal of DOX and Auxin and addition of Auxinole. Also at 56 hr, in ‘condition II’ cells RIF1 depletion was induced by addition of DOX and Auxin. 4 hr later (60 hr), both cell cultures were released from RO-3306. After 4 hr (64 hr), cells were seeded at 250/well in 6-well plates, then incubated for 7 days after which colonies were counted. **(C)** Cell cycle progression in HCT116 mAC-RIF1 cells during the procedure shown in B. Synchronisation was performed as in B in the presence of 1 μM Aphidicolin. Flow cytometry analysis was performed on a BD LSR Fortessa. The histograms show distribution of cellular DNA content at the time indicated. **(D)** Effect on colony formation rates of mAC-RIF1 depletion procedures combined with 1 μM Aphidicolin treatment, ‘condition I’ and ‘condition II’ cells treated according to the procedure shown in B. RIF1+ and RIF1-correspond to control mAC-RIF1 cells in which RIF1 was either expressed or depleted throughout the procedure. Values shown are normalised to the RIF1+ control. ***p<0.001.

Within the above synchronisation procedure, we either depleted RIF1 during the S phase Aphidicolin treatment period and re-expressed it for the recovery period (condition I, Fig. 2B), or else expressed RIF1 during the S phase treatment period and depleted it for the recovery period (condition II, Fig. 2B). We included control samples where RIF1 was either expressed or depleted throughout the entire experiment (Fig. 2D, RIF1+ and RIF1-). Cells depleted of RIF1 only during the replication stress treatment period (condition I) showed sensitivity similar to the RIF1-depleted (RIF1-) condition, displaying a surviving fraction of 27%. Cells depleted of RIF1 only during the recovery period (condition II) also showed high sensitivity, again similar to RIF1-with a surviving fraction of 18% (Fig. 2D). A repeat of this experiment (Fig. S1G) produced very similar results. These results indicate that molecular functions of RIF1 occurring both during and after a stressed S phase are important for survival. The cell death we observed in condition II did not simply reflect some effect of RO-3306, since cells expressing or lacking RIF1 showed no difference in RO-3306 sensitivity (data not shown).

We performed similar conditional depletion experiments in asynchronous cell populations (as outlined in S1H), and observed comparable results (Fig. S1I, S1J), in that depletion of RIF1 either during or after Aphidicolin treatment led to sensitivity.

To summarise, these results imply that RIF1 must be present during both treatment and recovery to protect cells from the effects of replication stress induced by Aphidicolin. The observation that RIF1 function remains important after a stressed replication period to promote cell survival is consistent with its highly dynamic pattern of localisation through late cell cycle stages (Fig. 1), consistent with RIF1 operating in chromosome maintenance processes occuring outside of S phase.

### Only the RIF1-Long splice isoform protects cells from replication stress

The RIF1 messenger RNA undergoes alternative splicing resulting in expression of ‘Long’ and ‘Short’ protein isoforms that we name RIF1-L and RIF1-S. RIF1-S lacks 26 amino acids corresponding to exon 31 (Fig. 3A, exon 31 shown in red; see Materials & Methods for Exon designation). We confirmed that transcripts encoding both the isoforms are widely expressed in various tissues *in vivo* (Table. S1). Both RIF1-L and RIF1-S transcripts are expressed in U2OS and HEK293 cell lines, and in human embryonic stem cells (not shown). Although they were reported as showing differential expression in cancer cells (Xu and Blackburn, 2004), distinct functions of the two isoforms have not previously been examined. We constructed cell lines expressing only mAC-RIF1-L or only mAC-RIF1-S, by inserting at the 3’ end of Exon 29 a ‘pre-spliced’ cDNA construct consisting of Exons 30-35 including or excluding Exon 31 (Fig. 3A). Clones were selected where both copies of the mAC-RIF1 gene contained the insertion. Western analysis confirmed that these mAC-RIF1-L and mAC-RIF1-S constructs encoded proteins expressed at levels similar to parental mAC-RIF1 (Fig. 3B, lanes 2 & 3).

**Figure 3:**
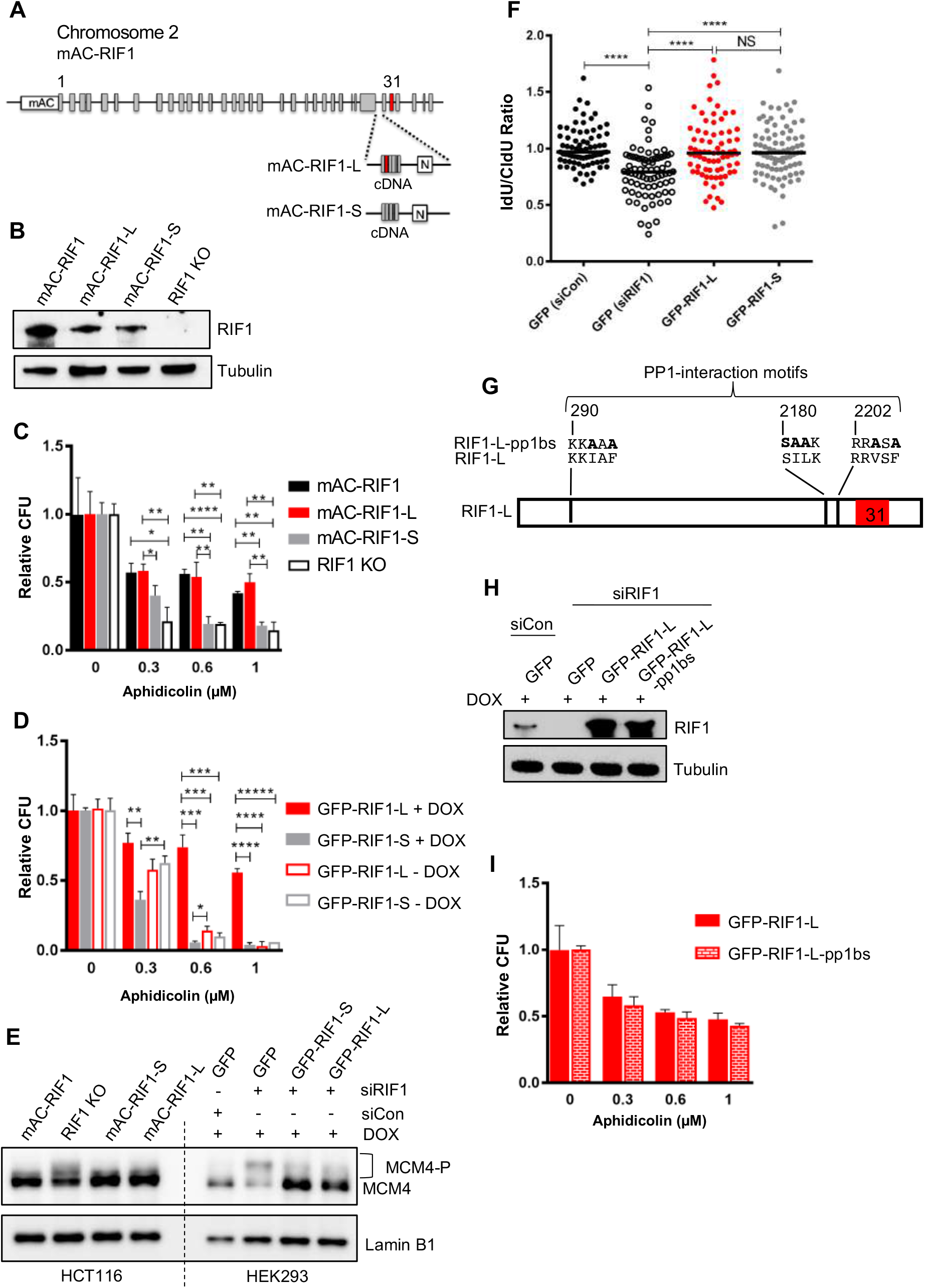
RIF1-L promotes resistance to Aphidicolin treatment but RIF1-S cannot. **(A)** Illustration of RIF1 exon structure and construction of HCT116 mAC-RIF1-L and mAC-RIF1-S cell lines. Constructs containing cDNA encoding the C-terminal portion of either RIF1-L or RIF1-S were inserted at the end of exon 29, by CRISPR-mediated integration of donor plasmids into the HCT116 mAC-RIF1 cell line. **(B)** Expression of RIF1 in HCT116 mAC-RIF1, mAC-RIF1-L, mAC-RIF1-S and RIF1 KO cell lines. Whole cell extracts were harvested for western blotting with anti-RIF1 antibody. Loading control shows tubulin. **(C)** CFA comparing resistance to Aphidicolin of mAC-RIF1(black bars), mAC-RIF1-L (red bars), mAC-RIF1-S (grey bars), and RIF1-KO (open bars) cell lines. *p<0.05; **p<0.01; ****p<0.0001. **(D)** CFA comparing effects of HEK293 GFP-RIF1-L (red bars) and GFP-RIF1-S (grey bars) expression on Aphidicolin resistance. **p<0.01; ***p<0.001; ****p<0.0001; *****p<0.00001. **(E)** Both RIF1-L and RIF1-S can counteract MCM4 hyperphosphorylation caused by depletion of endogenous RIF1. For HEK293 cells, 24 hr after transfection with either siControl or siRIF1, DOX was added to the culture medium. 24 hr later, chromatin-enriched protein fractions were prepared and analysed by western blotting with anti-MCM4 antibody. Hyperphosphorylated MCM4 protein (MCM4-P) is detected based on its retarded mobility as indicated by bracket. **(F)** DNA fiber assay. HEK293-derived cell lines were transfected with siCon or siRIF1 to deplete endogenous RIF1. The following day, expression of the stably integrated GFP, GFP-RIF1-L and GFP-RIF1-S was induced by addition of doxycycline (DOX). 2 days later cells were labeled with CldU and IdU, followed by a treatment with hydroxyurea (HU) 2 mM for 4 hours. DNA fiber assay was performed to assess nascent DNA degradation. Bar in the graph represents median value. 75 forks were analysed per sample and statistical significance was calculated using a Mann-Whitney test. NS; not significant; **** p ≤ 0.0001. **(G)** Illustration of RIF1 PP1-interaction motifs. To prevent PP1 interaction, critical residues in all three potential PP1 interaction motifs were replaced with alanine, creating a RIF1-pp1bs allele. **(H)** Expression of GFP-RIF1-pp1bs in HEK293 Flp-In T-Rex cells. 48 hr after transfection with siRIF1, DOX was added to the culture medium. After 24 hr, expression was assessed by western blotting with anti-RIF1 antibody. **(I)** CFA comparing effect of GFP-RIF1-L and GFP-RIF1-L-pp1bs expression on Aphidicolin resistance. The CFA was carried out as in Fig. 4C.

We found that while the mAC-RIF1-L cell line showed resistance to Aphidicolin very similar to that of the parent mAC-RIF1 cells (Fig. 3C, Fig. S2A, black and red bars), the mAC-RIF1-S isoform in contrast conferred little protection against drug, producing sensitivity similar to that of cells lacking RIF1 altogether (Fig. 3C, Fig. S2A, grey and open bars). This result suggested that only RIF1-L can protect cells from replication stress caused by Aphidicolin, and that RIF1-S is ineffective in this role.

To confirm this finding in a different cell line, we used HEK293-derived stable cell lines, containing siRIF1-resistant cDNA constructs encoding either GFP-RIF1-L or GFP-RIF1-S expressed under DOX control. Treatment of these cell lines with siRIF1, followed by DOX induction, allows replacement of endogenous RIF1 with either its Long or Short isoforms (Fig. S2B,C,D) (Hiraga et al., 2017). We found that also in this cell line RIF1-L was able to protect against Aphidicolin treatment, while RIF1-S could not (Fig. 3D, Fig. S2E, F, filled red and grey bars). Similarly, in the same system RIF1-L was able to protect cells from Hydroxyurea treatment while RIF1-S could not (Fig. S2G).

We considered molecular roles through which RIF1-L and RIF1-S might confer differing replication stress sensitivity. However, both isoforms appeared equally functional in the previously described mechanisms through which RIF1 affects DNA replication. In particular, the isoforms are equally effective in preventing hyperphosphorylation of the MCM complex (Fig. 3E & S2H) and protecting blocked replication forks (Fig. 3F). RIF1-S can therefore repress origin activation and protect nascent DNA, and yet cannot safeguard cells from Aphidicolin treatment, implying that responding to replication stress demands a further RIF1-mediated mechanism to promote cell survival, probably one that operates after the period of stress (Fig. 2) and that specifically requires RIF1-L (Fig. 3C, D).

In both its origin repression and nascent DNA protection functions, RIF1 acts as a PP1 substrate-targeting subunit (Hiraga et al., 2014, 2017; Kedziora et al., 2018). To further test for separability of the effect of RIF1 in replication stress survival from its known roles in replication control, we investigated whether PP1 interaction is essential for RIF1-L to protect against Aphidicolin treatment. We used the HEK293 cell line expressing a version of RIF1-L mutated at the PP1 interaction motifs to prevent PP1 interaction (Fig. 3G, H) (Hiraga et al., 2017). This ‘RIF1-L-pp1bs’ protein was almost as effective as wild-type RIF1-L in conferring resistance to Aphidicolin (Fig. 3I, Fig. S2I, J, indicating that RIF1-L acts largely independent of PP1 function in protecting cells from the effects of Aphidicolin. This independence from PP1 reinforces the evidence that the function of RIF1 in protecting from replication stress is distinct from its previously known roles in replication control, which do require PP1.

### RIF1-Long promotes 53BP1 nuclear body formation in G1 phase

We considered other routes through which RIF1 might promote survival after Aphidicolin treatment, focusing especially on events occurring after the replication stress period itself. UFBs form after replication stress, but we found no clear difference in localization of RIF1-L and RIF1-S to UFBs in mitotic cells (not shown). A further consequence of replication stress is the formation of large 53BP1 nuclear bodies in the subsequent G1 phase, which protect unreplicated DNA damaged by chromosome breakage at mitosis (Moreno et al., 2016; Lukas et al., 2011). We examined the formation of 53BP1 nuclear bodies 12 hr after removal of Aphidicolin in cells lacking RIF1-L, RIF1-S, or both RIF1 isoforms, limiting our analysis to G1 phase cells by counting only those that were cyclinA2-negative (Fig. S3A). The parental (mAC-RIF1) cell line showed an elevated fraction of cells with multiple large 53BP1 nuclear bodies (Fig. 4A top row, Fig. 4B, black bars and Fig. S3B top left panel), as expected. RIF1 was often co-localised with these bodies (Fig. 4A top row and Fig. 4C). In contrast, a reduced number of 53BP1 nuclear bodies was observed in RIF1-KO cells (Fig. 4A bottom row, Fig. 4B open bars, Fig. S3B bottom right panel), demonstrating that RIF1 contributes to the formation of 53BP1 bodies after replication stress. Examining the cell lines expressing only Long or Short RIF1 isoforms, we found that mAC-RIF1-L localised normally with 53BP1 bodies, and supported their formation at a near normal rate (Fig. 4A second row, 4B,C red bars and Fig. S3B top right panel). In contrast, the number of 53BP1 nuclear bodies formed in mAC-RIF1-S cells was similar to that in RIF1-KO cells, and mAC-RIF1-S showed reduced co-localisation with the 53BP1 bodies (Fig. 4A third row, 4B,C grey bars and Fig. S3B bottom left panel). The two RIF1 isoforms therefore differ in their effectiveness in promoting 53BP1 nuclear body formation following replication stress, with RIF1-L but not RIF1-S functional in this role. Since 53BP1 nuclear bodies are known to protect DNA damaged as a consequence of Aphidicolin treatment (Lukas et al., 2011), the defect in 53BP1 body formation in the absence of RIF1-L is likely to be a major factor in the replication stress sensitivity of RIF1-deficient cells, and may explain the isoform specificity of the replication stress protection function of RIF1.

**Figure 4:**
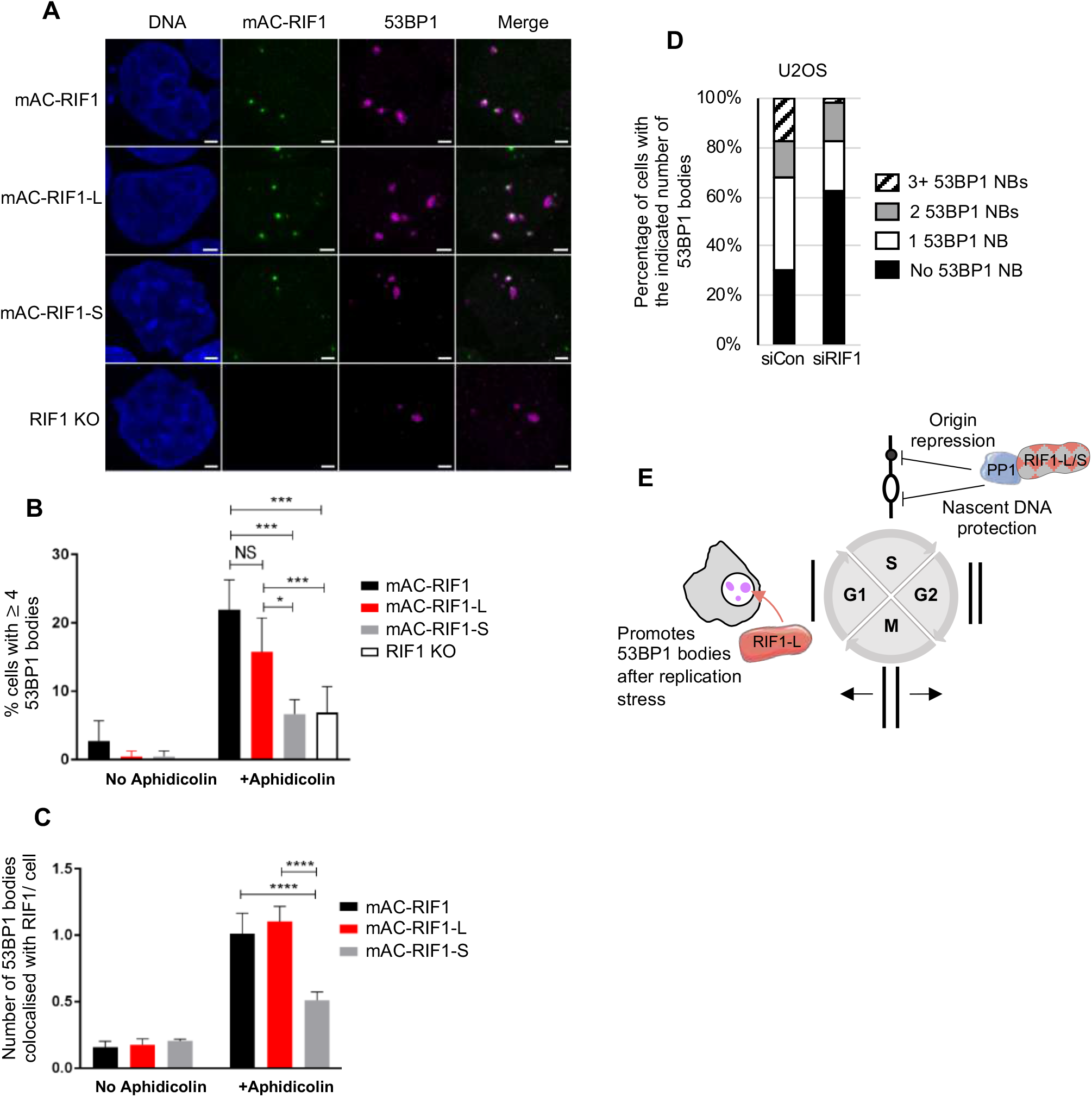
RIF1-L preferentially localises to 53BP1 protective nuclear bodies and promotes their formation. **(A)** Representative images showing 53BP1 nuclear bodies and RIF1 foci in HCT116 cell lines. Cells were treated with 1 μM Aphidicolin for 24 hr then released for 12 hr before fixing. Confocal microscopy was performed using an LSM880 + Airyscan (Zeiss). Scale bar = 2 μm. **(B)** Number of cells with 4 or more 53BP1 nuclear bodies in cells either untreated or treated with APH as in A. Plot shows averages and standard deviations from three experiments. NS; not significant, *p<0.05; ***p<0.001; ****p<0.0001. **(C)** Number of 53BP1 nuclear bodies colocalised with RIF1 foci per cells in untreated and APH-treated HCT116 cells. Averages and standard deviations taken from three independent experiments are shown. ****p<0.0001. **(D)** Distribution of 53BP1 NBs in U2OS cells treated as in A from one independent experiment. T-test (one-tailed, matched pairs), p = 0.046. **(E)** Illustration of RIF1-L and -S isoform functions.

To test in a different cell line whether RIF1 contributes to the number of 53BP1 nuclear bodies formed, we examined U2OS cells depleted of RIF1, 12 hr after release from a 24 hr Aphidicolin treatment (Fig. 4D, Fig, S3C). We found that RIF1 depletion caused a reproducible reduction in the number of nuclear bodies formed, similar to that observed in our HCT116-derived cell lines.

In investigating mechanisms through which RIF1 protects against interruption to replication, we tested RIF1 isoforms and found that RIF1-L is able to protect against replication stress while RIF1-S cannot. This deficiency of RIF1-S function was initially surprising, since RIF1-S appears competent to fulfil the known functions of RIF1 in DNA replication management—in particular RIF1-S is able to support replication licensing and control MCM phosphorylation (Fig. 3E & (Hiraga et al., 2017), and to protect against nascent DNA degradation (Fig. 3F & Fig. 4E). Consistently however, all of these known functions of RIF1 in replication control (promotion of replication licensing, control of MCM phosphorylation, and nascent DNA protection) depend on PP1 recruitment by RIF1 (Hiraga et al., 2014, 2017; Garzón et al., 2019); while we find that protection against replication stress does not require PP1 interaction (Fig. 3I, Fig. S3I,J), again suggesting that the role of RIF1 in protecting from replication stress might involve additional mechanisms.

Testing the effects of conditional depletion in synchronised cultures revealed that RIF1 is still needed after a period of replication stress to guard against toxicity, implying that protection from Aphidicolin involves a further function of RIF1. We therefore investigated whether RIF1 operates in post-S phase replication stress response pathways, a line of enquiry that revealed a role for RIF1-L in promoting the assembly of 53BP1 nuclear bodies (Fig. 4E). This specific function that we have discovered for RIF1-L may relate to a recently described role for RIF1 in ensuring the maturation and correct assembly of 53BP1 nuclear bodies following replication stress (Ochs et al., 2019), although the latter study did not measure number of nuclear bodies formed or test any differential contribution of the RIF1 isoforms. The finding that RIF1-L but not RIF1-S can function in promoting 53BP1 body formation potentially explains the specific requirement for RIF1-L in protecting against replication stress, since 53BP1 body assembly represents an important step in correct handling of stress-associated damage to enable ongoing proliferation. RIF1-L may act to directly promote 53BP1 body assembly, or possibly assist with the transit of damaged sites through to G1 phase to allow such protective bodies to form. Presently it is unclear how the apparently small difference of RIF1-L from the RIF1-S isoform (the inclusion of just 26 amino acids) specifically supports 53BP1 nuclear body formation. The findings described here do explain however that RIF1 contributes to recognising under-replicated and unrepaired sites for special protection and handling later in the chromosome cycle—in particular for delayed replication that guards against unscheduled recombinational repair to prevent the formation of pathological intermediates, as recently described (Spies et al., 2019). Overall, this study highlights the multifunctional role of RIF1 in ensuring chromosome maintenance to promote the survival and proliferation of cells after replication stress, emphasising the importance of RIF1 for determining response to replication-inhibiting chemotherapeutic drugs.

## Materials and Methods

### Cell lines

Stable HEK293 Flp-In T-Rex GFP and GFP-RIF1-S cell lines were as described (Hiraga et al., 2017; Escribano-Díaz et al., 2013). Constructed using the same procedure, described briefly below, were cell lines: HEK293 Flp-In T-Rex GFP-RIF1-L and HEK293 Flp-In T-Rex GFP-RIF1-L-pp1bs.

The HCT116 mAC-RIF1 cell line was constructed as as outlined below, using the system described (Natsume et al., 2016; Yesbolatova et al., 2019). HCT116 mAC-RIF1-L, HCT116 mAC-RIF1-S and HCT116 RIF1 KO cells were constructed as described below.

HCT116 mAC-RIF1 mCherry-PCNA was constructed by introducing the mCherry-PCNA construct under control of the EF1 alpha promoter using the piggyBac system(Yusa et al., 2011).

### HEK293 cell lines and culture conditions

HEK293-derived cell lines were cultivated in Dulbecco’s Modified Eagle’s Minimal medium supplemented with 10% foetal bovine serum (tetracycline-free), 100 U/ml penicillin, and 100 μg/ml streptomycin.

To construct cell lines, pOG44 (O’Gorman et al., 1991) and pcDNA5/FRT/TO-based plasmids carrying the GFP-RIF1-L or GFP-RIF1-L-pp1bs gene were mixed in 9:1 molar ratio and used for transfection of Flp-In T-Rex 293 cells (Invitrogen) with Lipofectamine 3000 (Invitrogen). Transfections and hygromycin B selection of stably transfected cells were performed as described by the manufacturer. Clones were tested for doxycycline-dependent induction of GFP fusion proteins by western blot and microscopy.

To assess the effect of ectopically expressing RIF1, cells were transfected with either control siRNA or siRNA against human RIF1. 2 days later, cells were split with addition of 1 μg/ml DOX then incubated for 24 hr to induce expression of GFP-RIF1 variant proteins. siRNA transfection was carried out using Lipofectamine RNAiMAX (Invitrogen) as described by the manufacturer. siRNA used were Human RIF1 siRNA (Dharmacon, D-027983-02: 5’-AGACGGTGCTCTATTGTTA-3’) and Control siRNA against Luciferase (Dharmacon, D-001100-01: 5’-CGTACGCGGAATACTTCGA-3’). Synonymous base mutations in the ectopically expressed GFP-RIF1 constructs make them resistant to siRNA targeted against endogenous RIF1 (Escribano-Díaz et al., 2013). RIF1 expression was assessed by western blot using RIF1 and GFP antibodies.

### HCT116 cell lines and culture conditions

HCT116-derived cells were cultivated in McCoys 5A medium supplemented with 2 mM L-glutamine, 10% foetal bovine serum (tetracycline-free), 100 U/ml penicillin, and 100 μg/ml streptomycin

#### HCT116 mAC-RIF1

To construct miniAID-mClover-fused RIF1 stable cell lines, HCT116 cells expressing the auxin-responsive F-box protein *Oryza sativa* TIR1 (OsTIR1) under the control of a Tet promoter were transfected using FuGENE HD (Promega) with a CRISPR/Cas9 plasmid targeting nearby the 1st ATG codon of the *RIF1* gene (5’-TCTCCAACAGCGGCGCGAGGggg-3’) together with a donor plasmid based on pMK345 (Yesbolatova et al., 2019), that contains a cassette (hygromycin resistance marker, selfcleaving peptide P2A, and mAID–mClover (Yesbolatova et al., 2019)) flanked by 500bp homology arms. Two days after transfection cells were diluted in 10 cm dishes, to which 100 μg/mL of Hygromycin B Gold (Invivogen) was added for selection. After 10-12 days, colonies were picked for further selection in a 96-well plate. Bi-allelic insertion of the donor sequence was checked by genomic PCR. Clones taken forward for analysis of RIF1 function were selected based on near endogenous RIF1 levels of RIF1 and efficient AID-mediated degradation of RIF1.

To induce degradation of miniAID-mClover-fused RIF1, OsTIR1 expression was first induced by 0.2 μg/ml DOX added to the culture medium, to produce a functional SCF (Skp1–Cullin–F-box) ubiquitin ligase that directs degradation of an AID-tagged protein (Nishimura et al., 2009; Natsume et al., 2016). After 24 hr, 10 μM Auxin (indole-3-acetic acid; IAA) was added to the culture medium to promote the interaction of mAC-RIF1 with SCF-OsTIR1, driving ubiquitination and mAC-RIF1 degradation. To suppress premature degradation of RIF1 in the presence of DOX, 100 μM of the TIR1 inhibitor Auxinole was added (Yesbolatova et al., 2019; Hayashi et al., 2012). In subsequent depletion experiments we used a regime in which DOX was first added in the presence of Auxinole, and then Auxinole was replaced with Auxin. Optimisation of DOX and Auxin concentrations is shown in Figure S1A and B. Unless otherwise stated, the above-mentioned drug concentrations were used throughout.

#### HCT116 mAC-RIF1-L/-S

HCT116 mAC-RIF1 cells were transfected with a CRISPR/Cas9 plasmid targeting the C-terminus of exon 29 of the *RIF1* gene (5’-CATCACCTGTTAATAAGGTAagg-3’) together with a donor plasmid, containing the cDNA of the C-terminal part of RIF1 (exons 30 to 35) with exon 31 (mAC-RIF1-L) or without (mAC-RIF1-S), followed by a Neomycin resistance marker (Figure 3A). Transfection and clonal selection were carried out as described above. Clones were tested for RIF1 expression by western blot.

#### HCT116 RIF1 KO

HCT116 cells were transfected with the same CRISPR/Cas9 plasmid that was used to construct HCT116 mAC-RIF1 cells, targeting near the 1^st^ ATG codon of the *RIF1* gene (sequence as above). Simultaneously transfected was a donor plasmid containing a hygromycin resistance marker flanked by 500bp homology arms. Transfection and clonal selection was carried out as described above. Clones were tested for loss of RIF1 expression by western blot.

#### PCR primers

The following PCR primers were used to construct pcDNA5/FRT/TO-GFP-RIF1-L:

SH572: 5’ – CTATGGAATTGAATGTAGGAAATGAAGCTAGC – 3’
SH593: 5’ – ACCGAGCTCGGATCGATCACCATGACGGCCAGGG – 3’
SH594: 5’ – GCCGCGGATCCGAATTCTAAATAGAATTTTCATGGGATGG – 3’
SH595: 5’ –
GCTACGTGATCCTGGGGACAGAAATCCTTTGGCTGAAGTGGTATTATGCTTAGAT
TGTGTAGTAGGAGAAG – 3’
SH596: 5’ –
TCCCCAGGATCACGTAGCCCTAAATTTAAGAGCTCAAAGAAGTGTTTAATTTCAG
AAATGGCCAAAG – 3’
SH597: 5’ – GATCAGTTATCTATGCGGCCG – 3’

The following PCR primers were used to amplify genomic DNA for the homology arms for the mAC-RIF1 donor plasmid pMK345:

HA1 For: 5’ - ccgggctgcaggaattcgatTAGGAGGGAGCGCGCCGCACGCGTG – 3’
HA1 Rev: 5’ – ggctttttcatggtggcgatCACCCTGAGGCCCGAACCGGAAGAG – 3’
HA2 For: 5’ - gctggtgcaggcgccggatccATGACGGCCAGGGGTCAGAGtCCCCTCGCGCC – 3’
HA2 Rev: 5’ – acggtatcgataagcttgatCTCTGGGTAGCCACATTTTCCCAAC – 3’

The following PCR primers were used to amplify genomic DNA for the homology arms for the RIF1 KO donor plasmid pMK194:

HA1 For: (see above)
HA3 Rev: 5’ - tcgctgcagcccgggggatcGGGGGCTCTGACCCCTGGCCGTCATGTCGG – 3’
HA4 For: 5’ – aagcttatcgataccgtcgaCTTTGGAAGACCCTTCTGCCTCCCATGGAG – 3’
HA2 Rev: (see above)

The following primers were used to amplify the C-terminal portion of either pcDNA5/FRT/TO-GFP-RIF1-L or pcDNA5/FRT/TO-GFP-RIF1 for the mAC-RIF1-L and mAC-RIF1-S donor plasmids:

5’ – AAATCTCATCACCTGTTAATAAG – 3’
5’ – acaagttaacaacaacaattCTAAATAGAATTTTCATGGGATGGT – 3’

The following primers were used to amplify the homology arms for the mAC-RIF1-L and mAC-RIF1-S donor plasmids:

5’ – ATGCAgagctcGAAACAGAGAATGAGGGCATAACTA – 3’
5’ – ATGCAggtaccATTCATTCAACAAACTATGTGCAAG – 3’

### Plasmids used for cell line constructions

RIF1 is encoded on chromosome 2. The RIF1 long variant cDNA (RIF1-L; NCBI RefSeq NM_018151.4) encodes a 2,472-amino acid protein, while the short variant (RIF1-S; RefSeq NM_001177663.1) lacks the 78-nucleotide stretch corresponding to exon 31 (Xu and Blackburn, 2004). We designate the exon containing the RIF1 ATG start codon as “exon 1”, so that our “exon 31” corresponds to “exon 32” of RefSeq NM_018151.4.

The GFP-RIF1 constructs used in this study are based on pcDNA5/FRT/TO-GFP-RIF1(Escribano-Díaz et al., 2013), which carries human RIF1-S cDNA with GFP fused at its N-terminus. To construct pcDNA5/FRT/TO-GFP-RIF1-L, a PCR fragment containing RIF1-S cDNA was amplified from pcDNA5/FRT/TO-GFP-RIF1 using primers SH593 and SH594, and cloned into pIRESpuro3 vector (linearised by EcoRV and EcoRI) using In-Fusion HD cloning system, to create plasmid pSH1009. The NheI-NotI fragment of the plasmid pSH1009 was replaced by two PCR fragments amplified by SH572 & SH595 and SH596 & SH597 respectively using In-Fusion HD system, to construct pSH1011 which has RIF1-L cDNA. The NheI-PspOMI fragment of pcDNA5/FRT/TO-GFP-RIF1 was replaced by NheI-NotI fragment of the pSH1011 plasmid to construct pcDNA5/FRT/TO-GFP-RIF1-L. Construction of a GFP-RIF1-S-pp1bs plasmid was previously described (Hiraga et al., 2017). The GFP-RIF1-L-pp1bs construct was made following a similar strategy.

The plasmid pX330-U6-chimeric_BB-CBh-hSPCas9 from Feng Zhang (Addgene, 42230) (Le Cong et al., 2013) was used to construct the CRISPR/Cas9 vector for the guide RNA (sequence as above) according to the protocol of Ran et al (Ran et al., 2013). Donor plasmids were based on pBluescript and constructed as described (Natsume et al., 2016; Yesbolatova et al., 2019). Primers for amplification of the homology arms and cDNA from RIF1-L and RIF1-S are listed under PCR primers.

### Protein extraction and western blotting

To prepare whole cell protein extracts, cells were trypsinised and washed with Dulbecco’s Phosphate-Buffered Saline (PBS) before treating with lysis buffer (10 mM Tris pH 7.5, 2 mM EDTA) containing a protease and phosphatase inhibitor cocktail (Roche). Chromatin-enriched protein fractions were prepared essentially as described (Mailand and Diffley, 2005). Protein concentrations were determined using the Bio-Rad RC-DC protein assay kit and equal amounts of total proteins were loaded in each lane. Proteins were transferred to PVDF membrane using the Trans-Blot Turbo Blotting System (Bio-Rad). Primary antibody incubation was overnight at 4°C (anti-RIF1, Bethyl Laboratories A300-568A; anti-MCM4, Abcam ab4459; anti-GFP, Abcam ab290; anti-MCM2 phospho-S53, Bethyl Laboratories A300-765A; anti-tubulin, Santa Cruz Biotechnology sc-53030). Proteins were detected by Clarity Western ECL blotting substrate (Bio-Rad) and the Bio-Rad Chemidoc Touch Imaging System.

### Colony formation assay (CFA)

In the case of HEK293-based cell lines, cells were seeded and transfected with 50 nM RIF1 siRNA using Lipofectamine RNAimax (Invitrogen) together with Optimem (Gibco). For HCT116-based cell lines, cells were seeded with 0.2 μg/ml DOX. After two days, cells were counted using the Invitrogen Countess II FL Automated Cell Counter and 250 cells added to each well of a 6-well plate, plated in triplicate. For HEK293-based cell lines, 1 μg/ml DOX was added to the culture medium to induce ectopic RIF1 expression whilst for HCT116-derived cell lines, 10 μM Auxin was added to the culture medium to degrade RIF1. Cells were incubated for 24 hr after which Aphidicolin (or Hydroxyurea) was added and cells incubated for a further 24 hr, before washing twice with PBS and replacement with the appropriate Aphidicolin-free medium. For HEK293-derived cell lines, 1 μg/ml supplementary DOX was re-added 72 hr after Aphidicolin removal and cells were then incubated for a further 4 days. In the case of HCT116-derived cell lines, after Aphidicolin removal, cells were incubated for 7 days. At the end of the incubation period, colonies consisting of more than 20 cells were counted directly using a Nikon Eclipse TS100 microscope, as in Buonomo et al (2009). Values were normalized to the 0 μM APH (DMSO) control. Statistical significance was calculated using a student’s T-test in Prism (Graphpad).

### Flow cytometry

To assess DNA content, cells were recovered by trypsinisation, then fixed with 70% ethanol. Cells were spun down and resuspended in 0.5 ml FxCyclePI/RNase staining solution (Molecular Probes, F10797) and incubated for 30 minutes at room temperature, protected from light. DNA content was analysed on a Becton Dickinson Fortessa analytical flow cytometer, and cell cycle distribution measured using FlowJo software. Doublet discrimination was performed by gating FSC-A against FSC-H.

To measure the mClover fluorescence of the mAC-RIF1 fusion protein, cells were recovered by trypsinsation and fixed with a 10% neutral buffered formalin solution (Sigma, HT-5012) for 30 minutes at 4°C, protected from light. Cells were washed with PBS/1% BSA before analysis on a Becton Dickinson Fortessa. mAC-RIF1 signal was measured using FlowJo software. Doublet discrimination was performed by gating FSC-A against FSC-H. An equal number of events are shown in each set of histogram plots.

### Cell cycle synchronisation

HCT116 cells were seeded in 12-well dishes and treated with 20 μM Lovastatin for 24 hr to induce G1 arrest (Rao et al., 1999). Cells were washed and medium containing 2 mM Mevalonic acid (MVA) added to induce release. 8 hr after release, 9 μM RO-3306 was added to hold cells at the G2/M boundary(Moghadam-Kamrani and Keyomarsi, 2008; Vassilev et al., 2006). After 28 hr, cells were washed and drug-free medium added allowing cells to enter mitosis. Flow cytometry was used to analyse synchronisation efficiency, and to establish and optimise the procedure for the experiment in Figure 3 based on assessment of cell cycle progression kinetics.

### DNA fiber assay

Cells were pulse-labelled with 50 μM CldU (20 min), followed by another pulse of 20 min of 250 μM IdU. After treatment with hydroxyurea (HU) 2 mM for 4 hours, cells were harvested and lysed on a microscope slide with spreading buffer (200 mM Tris pH 7.4, 50 mM EDTA, 0.5% SDS). Slides were tilted to allow the DNA suspension to run slowly and spread the fibers down the slide. Slides were fixed in cold (−20 °C) methanol-acetic acid (3:1) and DNA denatured in 2.5 M HCl at RT for 30 min. Slides were blocked and incubated with the following primary antibodies for 1 hour at RT in humidity chamber (anti-CldU, Abcam ab6326, 1:100; anti-IdU, BD 347580, 1:100; anti-ssDNA, Millipore MAB3034, 1:100). After washes with PBS, the slides were incubated with the following secondary antibodies (anti-rat IgG Alexa Fluor 594, Molecular Probes A-11007; anti-mouse IgG1 Alexa Fluor 488, Molecular Probes A-21121; anti-mouse IgG2a Alexa Fluor 350, Molecular Probes A-21130). Slides were air-dried and mounted with Prolong (Invitrogen). Samples were imaged under a Zeiss Axio Imager and analysed using ImageJ. CldU and IdU tract lengths were measured in double-labelled forks and the IdU/CldU ratio was used as an indicator of nascent DNA degradation.

### Immunofluorescence microscopy

HCT116 cells were cultured in a glass-bottomed dish (MatTek). Cells were fixed with 4% paraformaldehyde/PBS, for 15 min at room temperature. Cells were antibody stained with primary antibodies (anti-53BP1, Santa Cruz Biotechnology sc-22760; anti-FANCD2, Novus Biologicals NB100-182; anti-BLM, Santa Cruz Biotechnology sc-7790) and secondary antibodies (anti-rabbit Alexa Fluor 594, Thermo Fisher Scientific A-11037; anti-goat Alexa Fluor 594, Thermo Fisher Scientific A-11058; GFP-Booster ATTO 488, ChromoTek gba488) according to the manufacturer’s guidelines. Microscopy was performed using a DeltaVision Personal (GE) microscope with a x60 objective. 41 Z-stacks of 0.3 μM were taken and deconvolution was performed using the SoftWoRx software (DeltaVision). Maximum intensity projections (MIP) were created using Volocity software (PerkinElmer).

### Live cell imaging

HCT116 cells were cultured in a glass-bottomed dish (MatTek) containing the medium without phenol red. To visualize nuclei in live cells, 0.5 μM SiR-DNA (Spirochrome) was added to the medium before observation. Live cell imaging was performed using a DeltaVision microscope equipped with an incubation chamber and a CO_2_ supply (GE healthcare Life Sciences). Image analysis and quantification were performed using the Volocity software (PerkinElmer). The half-life of mClover signal was calculated using the Prism software (GraphPad).

### Confocal microscopy

HCT116 cells and U2OS cells were cultured in ibiTreat μ-slide 8 well dishes (Ibidi) at 37 °C with 5% CO_2_. Cells were treated with 1 μM Aphidicolin for 24 hr after which Aphidicolin was removed and cells were incubated for a further 12 hr. Cells were fixed with either a 10% neutral buffered formalin solution (Sigma, HT-5012) for 10 minutes at room temperature or with 100% methanol for 15 minutes at −20 °C. Blocking was for 30 minutes with PBST/1%BSA at 4 °C after which antibody staining was performed (anti-cyclin A2-Alexa Fluor 555, Abcam ab217731; anti-53BP1, Novus Biologicals NB100-94 self-conjugated to Alexa Fluor 647, Expedeon 336-0030). Confocal microscopy was performed using an LSM880 + Airyscan (Zeiss). x63 magnification was used and Z-stacks were imaged (40 slices). Images were processed first to an airyscan image and then to a maximum intensity projection (MIP) using ZEN Black (Zeiss). Cyclin A2 staining was used to identify G1 cells (Cylin A2 negative = G1 cells). Both image analysis (colocalization) and quantification were performed using CellProfiler (Broad Institute) with a custom-made pipeline. The ‘IdentifyPrimaryObjects’ module was used to identify individual cells (DAPI positive), the G1 cell population (CyclinA2 negative), mAC-RIF1 foci (mClover positive) and 53BP1 foci (Alexa Fluor 647 positive). The ‘RelateObjects’ module was used to count the number of mAC-RIF1 and 53BP1 foci in a cell and to assess colocalization of 53BP1 foci with mAC-RIF1 foci. Statistical significance was calculated using a one-way ANOVA using Prism (Graphpad).

### cDNA analysis of RIF1 splice variants

A human cDNA panel covering 48 major tissues was obtained from Insight Biotechnology (Origene HMRT104). Two RIF1 splicing variants were amplified by competitive PCR using a single primer pair (LW030 and LW031). The PCR product was analysed on a 2%agarose gel (3:1 mix of low-melting temperature agarose and normal agarose). Bands were visualised by staining with SYBR Green I dye, and fluorescent images were captured using Bio-Rad ChemiDoc Touch system. Intensities of bands specific to RIF1-L, RIF1-S and an additional minor band caused by heteroduplex formation, were measured using ImageLab software (Bio-Rad). The relative abundance of RIF-L and RIF1-S were calculated based on the band intensities, with 50% of the heteroduplex band intensity assigned to RIF1-L and 50% assigned to RIF1-S.

Primers used

- LW030 (5’ GTCTCCTTTGGCTTCTCCGT 3’)
- LW031 (5’ GATGTCAACTGGTGCCACAC 3’)

## Acknowledgements

Thanks to members of the Aberdeen Chromosome Stability Group and to Kevin Hiom for helpful comments. We thank Raif Yuecel and his team at the Iain Fraser Cytometry Centre for assistance, and Kevin McKenzie and his team at the Microscopy and Histology Core Facility. Work at the University of Aberdeen was supported by Cancer Research UK Studentship Award C1445/A20596 and CRUK Programme Award C1445/A19059. Work at the National Institute of Genetics, Mishima was supported by JSPS KAKENHI Grants Numbers 17K15068, 18H02170 and 18H04719, and by research grants from the Daiichi Snakyo Foundation of Life Science, and the Takeda Science Foundation. Collaboration was supported by NIG-JOINT (98I2019) and by a 2017 JSPS Summer Programme Fellowship.

## Author Contributions

LPW, SH, TN, MK and ADD conceived and designed the experiments. LPW, SH, TN, YS, JG and MK performed experiments. LPW, SH and TN analysed the data. LPW, SH and ADD wrote the manuscript.

---

**Supplementary Figure 1: Characterisation and optimisation of mAC-RIF1 depletion (A)** Optimisation of DOX concentration for TIR1 induction in HCT116 mAC-RIF1 cell line. Cells were treated with a range of DOX concentrations as indicated and mClover signal was analysed by flow cytometry, establishing that 0.2 μg/ml DOX is sufficient for degradation. **(B)** Optimisation of Auxin for SCF-OsTIR1-mediated RIF1 depletion in HCT116 mAC-RIF1 cell line. Cells were treated with a range of Auxin concentrations as indicated and mClover signal was analysed by flow cytometry, establishing that 10 μM Auxin is sufficient for degradation. **(C)** mAC-RIF1 depletion analysed by flow cytometry. Cells were treated with 2 μg/ml DOX and Auxinole (to suppress reduction in mAC-RIF1 flourescence signal from OsTIR1 induction by treatment with DOX alone) for 24 hr (plot 4 from top), then medium was replaced with 2 μg/ml DOX and 500 μM Auxin for the indicated time periods (plots 5-8). Plots 1-3 show mClover signal in untagged cells and in cells treated with no drug or DOX only. Samples were taken at the indicated time points and mClover signal analysed by flow cyometry. mAC-RIF1 depletion was largely complete after 3 hr of Auxin treatment, as assessed by comparison with a DOX + Auxin 24 hr sample (seventh and eighth plots). **(D)** Testing re-expression of mAC-RIF1 following a period of depletion. RIF1 was first depleted by adding DOX and Auxin for 24 hr. Cells were washed three times after which Auxinole-containing medium was added. Samples were taken at the indicated time points and mClover signal analysed by flow cytometry. Based on the kinetics of RIF1 depletion and reexpression, a 4 hr window was allowed for complete depletion or re-expression of RIF1. **(E)** CFA testing Aphidicolin sensitivity of HCT116 mAC-RIF1 cells depleted for RIF1 (by DOX and Auxin) and HCT116 RIF1 KO cells. Cells were seeded at low density and HCT116 mAC-RIF1 cells treated with 0.2 μM DOX and 10 μM Auxin for 24 hr before treatment with Aphidicolin concentrations indicated. **(F)** Synchronisation of HCT116 mAC-RIF1 cells. Cells were arrested at G1 phase with Lovastatin for 24 hr (‘G1 arrest’) then released with Mevalonic acid (no Aphidicolin). After 8 hr, RO-3306 was added to arrest cells in G2 phase. 19 hr later (‘G2 arrest’), cells were released by washing out the RO-3306, and sampled 4 hr later. Samples were taken at the indicated points through the experiment and DNA stained with PI prior to flow cytometry analysis. **(G)** Effect on colony formation rates of mAC-RIF1 depletion during or after 1 μM Aphidicolin treatment, treated according to ‘condition I’ or ‘condition II’ procedures shown in Fig. 2B. RIF1+ and RIF1-correspond to control mAC-RIF1 cells with RIF1 either expressed or depleted throughout the experiment. Experiment represents a duplicate of that in Fig. 2D. Values were normalised to the RIF1+ condition. *p<0.05; ***p<0.001; ****p<0.0001. **(H)** Outline of procedure for testing effect of RIF1 depletion either during or after Aphidicolin treatment, in unsynchonised cultures. Cells were seeded at 250 cells/well in 6-well plates in the presence of DOX and Auxinole. On day 2, Aphidicolin was added, and then removed on day 3. Cells were incubated for 7 days after which colonies were counted using a Nikon Eclipse TS100 microscope. For condition III (treatment RIF1- / recovery RIF1+), RIF1 was depleted during the drug treatment period, with DOX and Auxin added to the medium 3 hr before Aphidicolin was added. 3 hr before the end of the drug treatment, RIF1 was re-expressed by replacing medium with medium containing Auxinole. For condition IV (treatment RIF1+ / recovery RIF1-), DOX and Auxinole were maintained in the medium until 3 hr before the end of the drug treatment when it was replaced with medium containing DOX and Auxin to induce RIF1 depletion. **(I)** Results of experiment in Fig. S1H testing whether RIF1 is needed during or after drug treatment to protect cells from replication stress. HCT116 mAC-RIF1 cells were depleted of RIF1 either during the 24 hr Aphidicolin treatment (condition III, horizonal lined bars) or during the recovery period (condition IV, hatch pattern bars). For the ‘RIF1-’ condition (open bars), RIF1 was depleted by DOX and Auxin addition 24 hr before drug treatment. *p<0.05; **p<0.01; ***p<0.001. **(J)** Duplicate experiment of that shown in part H & I, except that a window of 4 hr was used for depletion and re-expression of RIF1. That is, for condition III DOX and Auxin added to the medium 4 hr before Aphidicolin was addition, then replaced with Auxinole medium 4 hr before the removal of Aphidicolin; while for condition IV medium contained DOX and Auxinole until 4 hr before the end of the drug treatment, then replaced with medium containing DOX and Auxin. *p<0.05; **p<0.01; ***p<0.001.

**Supplementary Figure 2: RIF1-L promotes resistance to Aphidicolin treatment but RIF1-S cannot (A)** CFA comparing resistance of mAC-RIF1(black bars), mAC-RIF1-L (red bars) mAC-RIF1-S (grey bars), and RIF1-KO (open bars) cell lines to treatment with different Aphidicolin concentrations. *p<0.05; **p<0.01. Experiment represents a duplicate of that in Fig. 3C. **(B)** Illustration of DOX-controlled GFP-RIF1-L and GFP-RIF1-S cDNA constructs. Following siRNA-mediated depletion of endogenous RIF1, splice variant-specific expression of RIF1 is induced by the addition of 1 μg/ml DOX. **(C)** Expression of endogenous RIF1, or GFP-RIF1-L and GFP-RIF1-S, in cells treated with siControl or siRIF1. DOX was added to the culture medium 24 hr before whole cell extracts were harvested for western blotting. The same protein samples are shown probed with anti-RIF1 (top), anti-GFP (middle), or loading control anti-tubulin (bottom). **(D)** Modified CFA assay. On Day 1, GFP-RIF1-L and GFP-RIF1-S cells were transfected with siRIF1 or non-targeting control siRNA. On day 3, cells were seeded with DOX to induce transcription of siRNA-resistant GFP-RIF1-L or GFP-RIF1-S constructs. On day 4, Aphidicolin was added then removed on day 5. Cells were incubated for a further 7 days then colonies counted on day 12. **(E)** CFA comparing effects of HEK293 GFP-RIF1-L (red bars) and GFP-RIF1-S (grey bars) expression on Aphidicolin resistance. Experiment was carried out as in S2D, with Aphidicolin treatment at concentrations indicated. *p<0.05; **p<0.01; ***p<0.001; ****p<0.0001. Experiment represents a duplicate of that in Fig. 3D. **(F)** CFA comparing effects of HE293 GFP (siControl) (black spotted bars), HEK293 GFP-RIF1-L (siRIF1) (red bars) and GFP-RIF1-S (siRIF1) (grey bars) expression on Aphidicolin resistance. Experiment was carried out as in S2D, with Aphidicolin treatment at concentrations indicated. **p<0.01; ***p<0.001. **(G)** CFA comparing effects of Hydroxyurea on HEK293-derived cells expressing only GFP-RIF1-L (red bars) or only GFP-RIF1-S (grey bars) in place of endogenous RIF1. Experiment was carried out as in S2D, but replacing Aphidicolin with Hydroxyurea treatment at concentrations indicated. **p<0.01; ***p<0.001; ****p<0.0001. **(H)** RIF1-L and RIF1-S can counteract MCM2 hyperphosphorylation caused by depletion of endogenous RIF1. For HEK293 cells, 24 hr after transfection with either siControl or siRIF1, DOX was added to the culture medium. 24 hr later, chromatin-enriched protein fractions were prepared and analysed by western blotting with anti-p-S53-MCM2 antibody. Hyperphosphorylated MCM2 shows a specific p-S53-MCM2 band as indicated by the arrow, with the phosphorylated form running faster. **(I)** CFA comparing effect of GFP-RIF1-L and GFP-RIF1-L-pp1bs expression on Aphidicolin resistance. The CFA was carried out as in Fig. S2D. Experiment represents a duplicate of that in Fig. 3I. **(J)** CFA comparing effect of GFP-RIF1-L and GFP-RIF1-L-pp1bs expression on Aphidicolin resistance. The CFA was carried out as in Fig. S2D. Experiment represents a duplicate of that in Fig. 3I. **p<0.01.

**Supplementary Figure 3: Distribution of 53BP1 nuclear bodies after Aphidicolin treatment (A)** Representative images of CyclinA2 staining in experiment shown in Fig. 4A. Circled cell correspond to those in the panels of Fig. 4A. Scale bar = 5 μM. **(B)** Distribution of the number of 53BP1 nuclear bodies per cell as a % of the cell population, from same cultures as in Figure 4A. Average values and standard deviations from three independent experiments are shown. **(C).** Distribution of 53BP1 NBs in U2OS cells treated as in Fig. 4A. Two independent experiments are shown. T-test (one-tailed, matched pairs), p = 0.046.

**Supplementary Table 1: RIF1 splice variants are expressed *in vivo*** RIF1 splice variants were amplified from a human cDNA panel covering 48 major tissues. The PCR products were analysed on a 2% agarose gel and the relative abundance of RIF1-L and RIF1-S were calculated based on the band intensities using ImageLab Software (Bio-Rad).

**Supplementary Video 1: mClover-tagged RIF1 from early to late S phase** Time-lapse imaging video of unsynchronised HCT116 mAC-RIF1 mCherry-PCNA cells transitioning from early to late S phase. Blue represents DNA (DAPI stain), red represents PCNA (mCherry-PCNA) and green represents mAC-RIF1 (mClover).

**Supplementary Video 2: mClover-tagged RIF1 from late S to the following G1 phase** Time-lapse imaging video of unsynchronised HCT116 mAC-RIF1 mCherry-PCNA cells transitioning from late S to the following G1 phase. Blue represents DNA (DAPI stain), red represents PCNA (mCherry-PCNA) and green represents mAC-RIF1 (mClover).

