## Supplemental figures and table for "RIF1-Long promotes G1 phase 53BP1 nuclear bodies to protect against replication stress"

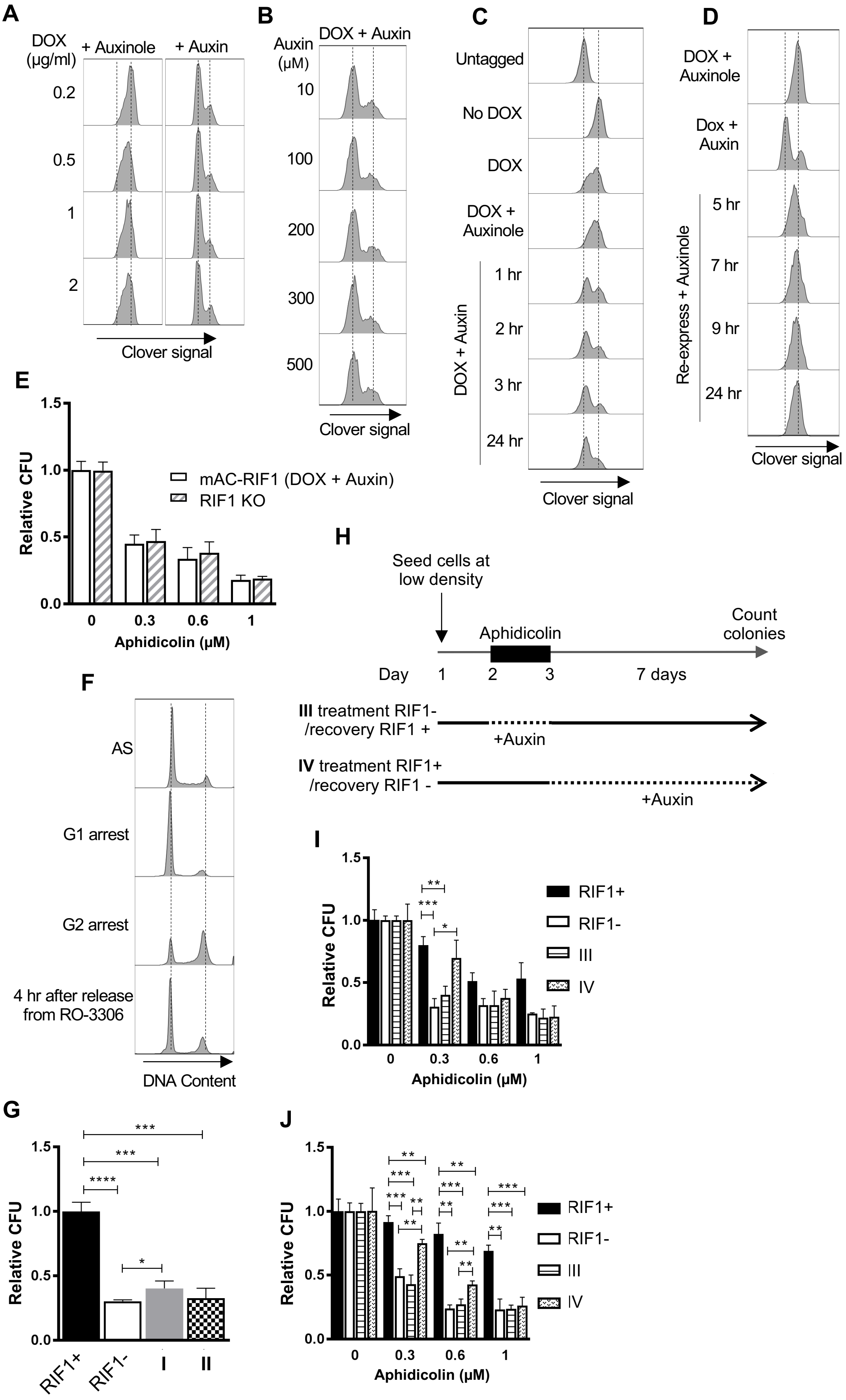


Watts_SupFig1

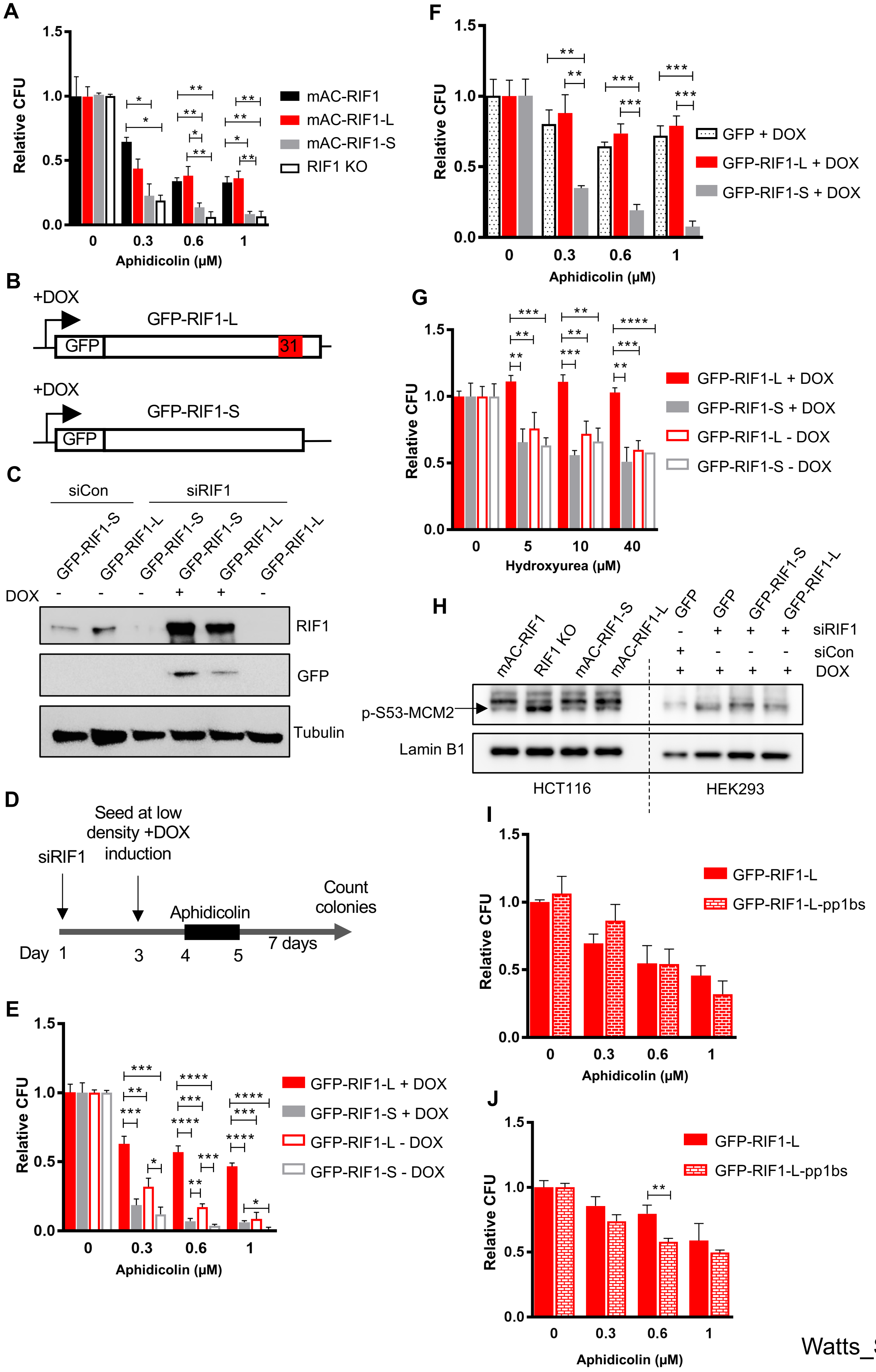


Watts_SupFig2

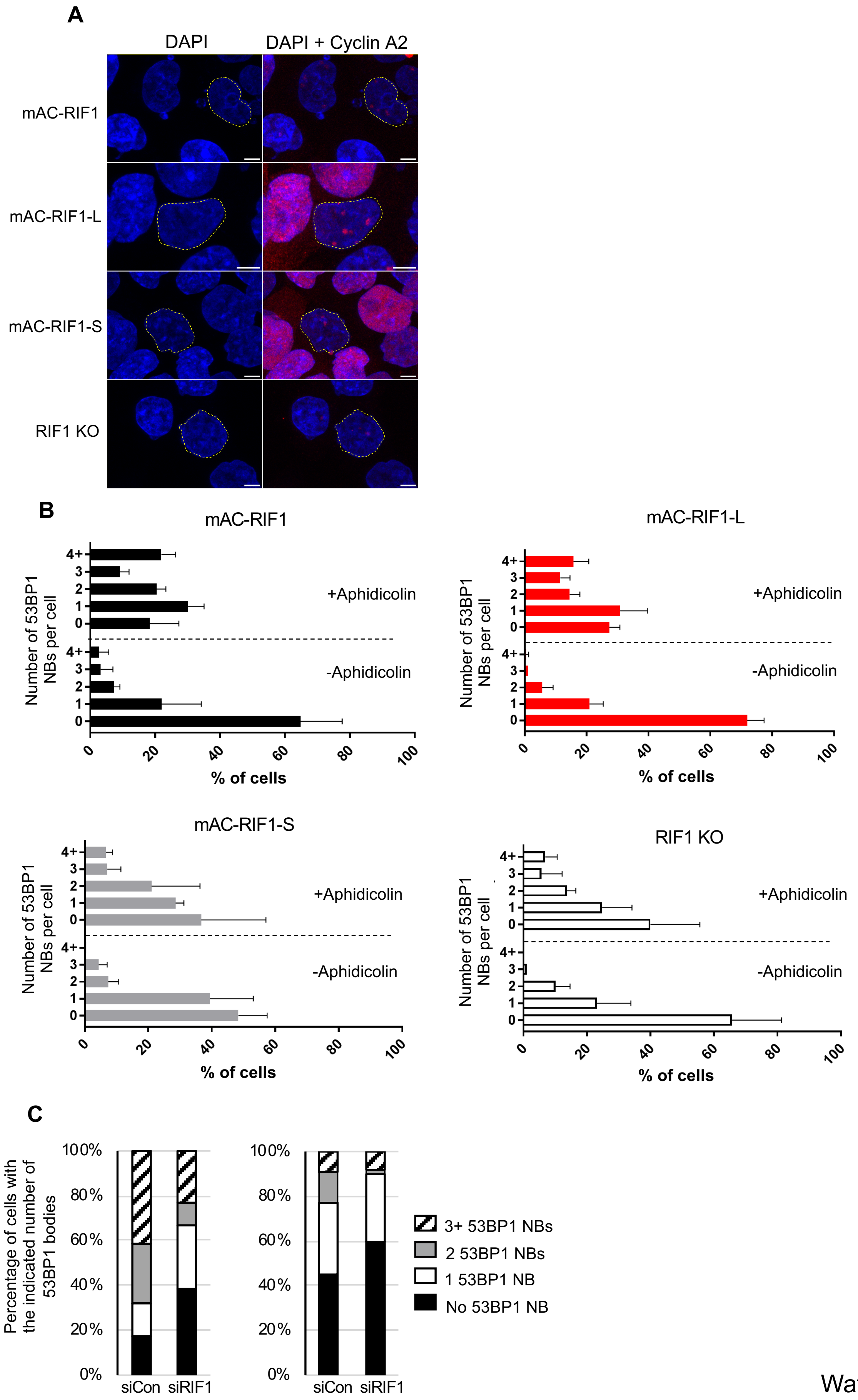


Watts_SupFig3

| Tissue | % L | % S |
|---|---|---|
| Adrenal Gland | 63 | 37 |
| Bone marrow | 70 | 30 |
| Brain | 85 | 15 |
| Bronchus | 75 | 25 |
| Cervix | 75 | 25 |
| Colon | 64 | 36 |
| Descending part of duodenum | 72 | 28 |
| Epididymis | 74 | 26 |
| Esophagus | 71 | 29 |
| Heart | 83 | 17 |
| Heart, Ventricle (left) | 80 | 20 |
| Heart, Ventricle (right) | 76 | 24 |
| Intestine (small) | 75 | 25 |
| Kidney | 67 | 33 |
| Larynx | 65 | 35 |
| Liver | ND | ND |
| Lung | 77 | 23 |
| Lymph node | 69 | 31 |
| Lymphocytes (peripheral blood) | 68 | 32 |
| Mammary gland | 100 | 0 |
| Muscle | 62 | 8 |
| Nasal mucosa | 84 | 16 |
| Optic nerve | 85 | 15 |
| Ovary | 63 | 37 |
| Oviduct | 77 | 23 |
| Pancreas | 76 | 24 |
| Penis | 0 | 100 |
| Pericardium | 87 | 13 |
| Pituitary | 83 | 17 |
| Placenta | 63 | 37 |
| Prostate | 69 | 31 |
| Retina | 78 | 22 |
| Salivary gland | 78 | 22 |
| Skin | 64 | 36 |
| Spinal cord | 75 | 25 |
| Spleen | 70 | 30 |
| Stomach | ND | ND |
| Testis | 44 | 56 |
| Thymus | 60 | 40 |
| Thyroid | 82 | 18 |
| Tongue | ND | ND |
| Tonsil | 70 | 30 |
| Trachea | ND | ND |
| Ureter | 71 | 29 |
| Urinary bladder | 77 | 23 |
| Uterus | 81 | 19 |
| Uvula | 64 | 36 |
| Vagina | 78 | 22 |
